# Genotype-specific variation in seasonal body condition at a large-effect maturation locus

**DOI:** 10.1101/2023.01.15.524064

**Authors:** Andrew H. House, Paul V. Debes, Johanna Kurko, Jaakko Erkinaro, Craig R. Primmer

## Abstract

1. Organisms utilize varying lipid resource allocation strategies as a means to survive seasonal environmental changes and life-history stage transitions. In Atlantic salmon, a certain lipid threshold is needed to initiate sexual maturation. Because of this, an individual’s maturation schedule may be affected by changes in temperature and food availability across the seasons that create natural fluctuations of lipid reserves.
2. Recent studies have found a genome region, including the gene *vgll3,* that explains a large proportion of variation for size and age at maturity*. Vgll3* encodes a transcription co-factor that acts as an inhibitor of adipogenesis in mice and also affects condition factor and other phenotypes in juvenile salmon. However, even with many studies investigating varying temperature effects, there is a lack of temporal studies examining the effects of seasonality on such phenotypes, nor have the effects of *vgll3* genotype on condition factor and maturation in different temperatures at different life stages.
3. Here, we investigate the influence of different larval and juvenile incubation temperatures, *vgll3* genotype and their interactions on juvenile salmon phenotypes including body condition, and sexual maturation rate. We reared Atlantic salmon for 2 years in varying temperatures with an average 1.76 °C difference between warm and cold treatments in four different larval-juvenile phase treatment groups (Warm-Warm, Warm-Cold, Cold-Warm, and Cold-Cold) until the first occurrence of maturation in males.
4. We found no effect of larval temperature on the measured phenotypes or maturation rate, suggesting the occurrence of growth compensation over the course of the experiment. Agreeing with previous studies, an increased maturation rate was observed in individuals of the warm juvenile temperature treatment.
5. In addition, we observed differences in condition factor associated with *vgll3* genotype, whereby *vgll3*EE* individuals (the genotype associated with early maturation) had a less variable condition factor across the seasons compared to the *vgll3*LL* (associated with late maturation) individuals.
6. This result suggests a *vgll3* influence on resource acquisition and allocation strategies, possibly linked with the early maturation process, with individuals carrying the early maturation *vgll3* genotype having a higher early maturation rate and a higher condition factor in the spring.

## Introduction

Resource acquisition and allocation strategies are important for enabling organisms to respond to environmental fluctuations (Mogensen and Post 2012). In ectotherms, temperature is an important abiotic factor that can directly influence food availability but also growth and metabolism, and thereby development rate (Castañeda, Lardies, and Bozinovic 2004; Finstad and Jonsson 2012; Jonsson, Jonsson, and Finstad 2014). Because of the many influences temperature has it also can affect an individual’s ability to use or store acquired energy (Geissinger et al. 2021; Mogensen and Post 2012; Post and Parkinson 2001). The amount and usage of the stored energy reserves plays a role in initiating and progressing developmental processes, such as life-history stage transitions associated with smoltification (i.e., acquiring seawater tolerance) and sexual maturation (Jonsson and Jonsson 2005; reviewed in Wang, Hung, and Randall 2006). Therefore, understanding the processes that determine the allocation of acquired energy will contribute to understanding variation in initiation and progression of major life-history transitions (Post and Parkinson 2001; Rowe, Thorpe, and Shanks 1991).

Lipid allocation patterns can affect the probability of survival to reproductive age as well as the probability of maturation at a given age (Post and Parkinson 2001). Condition factor, defined as the relative weight of an individual given its length, provides an indication of the relative level of lipid stores in Atlantic salmon during the freshwater stage (Herbinger and Friars 1991; Sutton, Bult, and Haedrich 2000). It has also been applied as a proxy for lipid reserve levels in a range of taxa including birds (Balbontín et al. 2012), amphibians (Cogălniceanu et al. 2021), mammals (Bright Ross et al. 2021), and fishes (Haraldstad et al. 2018; Mozsár et al. 2015; Sutton, Bult, and Haedrich 2000). Lipid reserves have been found to be especially important for Atlantic salmon due to their need to reach a certain threshold of lipid reserves in order to initiate maturation (Rowe, Thorpe, and Shanks 1991; Rowe and Thorpe 1990a; 1990b) which may be affected by variation for replenishing reserves across seasonal changes, including periods of low temperatures and food availability (Gurney et al. 2003; Mogensen and Post 2012). For example, it has been shown that maturing males replenish and build up stores faster than individuals delaying maturation in the spring prior to maturation capability (Kadri et al. 1996; Rowe, Thorpe, and Shanks 1991).

Atlantic salmon is an excellent organism to understand energy allocation and effects of environmental variation at different life stages. The propensity to mature early has been associated with prior condition factor (Debes et al. 2021; Herbinger and Friars 1992; Rowe, Thorpe, and Shanks 1991). Further, the genetic basis of age at maturity in salmon has been well characterized, with a single genome region, including the *vgll3* gene, explaining 39% of the variation in the age at maturity (Barson et al. 2015; Czorlich et al. 2018). *Vgll3* encodes a transcription cofactor and has been associated with adipocytes differentiation in mice (Halperin et al., 2013) and recently was also found to play a role in mediating maturation timing via condition factor in salmon (Debes et al. 2021). However, the effects of *vgll3* genotype on condition factor and maturation in different temperatures at different life stages has not been investigated.

To address this knowledge gap, we reared Atlantic salmon with different *vgll3* genotypes from fertilization for two years in four different temperature treatment combinations: warmer or colder (2°C difference) during the embryonic and larval endogenous feeding phase (fertilization to first feeding, hereafter ‘larval’) and warmer or colder during the externally feeding juvenile phase (hereafter ‘juvenile’). This enabled us to study the relative effects of environmental temperature during larval and juvenile rearing on resource allocation relevant phenotypes, such as growth, body condition, maturation rate and size at maturity. We also investigated whether genetic effects or interactions with the environmental effects (GxE) exist in order to understand seasonal energy allocation and its effect on the maturation process.

## Methods

### Salmon rearing and measurement

The Atlantic salmon used in this study derived from a first-generation hatchery stock maintained by the Natural Resources Institute Finland (62°24ʹ50″N, 025°57ʹ15″E, Laukaa, Finland), which originate from the River Neva, Russia. Fertilization took place in late October 2017 when unrelated parents with homozygous *vgll3* genotypes were crossed as six 2 × 2 factorials. Each factorial included a *vgll3*EE* male and female and a *vgll3*LL* male and female, where *E* and *L* refer to the alleles previously associated with **e**arlier or **l**ater maturation, respectively (Barson et al. 2015), thus yielding four reciprocal same-*vgll3*-genotype offspring families in each 2×2 factorial (EE, 2EL, LL). Eggs of the 24 families were divided into four batches and incubated in each of two vertical incubators at each of two temperatures (2°C difference, hereafter referred to as the warm and cold larval treatments), i.e., with two replicates per family and temperature treatment. A water temperature difference of 2°C was maintained by using a combination of water chillers and room heating. At first feeding, juveniles from the two replicates were pooled and transported to the Lammi Biological Station (61°04′45′′N, 025°00′40′′E, Lammi, Finland) on 10.03.2018 and 24.04.2018 for the warm- and cold-larval treatments, respectively. Half of the individuals of each larval temperature treatment were placed into the same temperature treatment for the juvenile phase (warm and cold juvenile treatments; maintaining a 2°C difference), and the other half of the individuals were transferred to the opposite temperature treatment, thus resulting in a total of four different larval phase-juvenile phase temperature treatment groups, Warm-Warm (WW), Warm-Cold (WC) , Cold-Warm (CW), and Cold-Cold (CC) as shown in Figure 1A. Each treatment was replicated in five flow-through circular tanks (diameter 90 cm), and juveniles of each family were allocated to their respective replicate treatment tanks in roughly equal numbers and subsequently reared under a controlled photoperiod set to the local latitude *(coordinates). Water was sourced from a nearby lake, Lake Pääjärvi, and thus followed the natural annual water temperature cycle with the cold and warm water treatment maintained via a heat-exchange system ranging from 1.30-18.53°C and 1.35-19.04°C, respectively, with an average difference of 1.76°C (Figure 1B). Fish were fed *ad libitum* with commercial fish food, the pellet size of which matched the requirements set by the size distribution of the individuals (Raisio Baltic Blend; Raisio Oy) for the duration of the experiment. Wet mass (± 0.01 g) and fork length (± 1 mm) were measured, and a fin clip sampled for genetic analysis, for a sub-set of individuals at five timepoints (464-580 per time point), the first of which was eight months post-fertilization, and the last when the experiment was terminated at 24 months post-fertilization (Figure 1B). Sex was determined phenotypically following dissection, and maturation status was also assessed at the last two measuring timepoints through dissection and gonad assessment. In October 2018, an accident during cleaning resulted in the loss of one tank per treatment group, resulting in four replicate tanks per treatment group for sampling time points 2-5. Following DNA extraction, samples were genotyped with 141 SNPs and a sexing marker (Aykanat et al. 2016) to determine *vgll3* genotype and assign family of origin as outlined in Debes et al. (2021).

**Figure 1:**
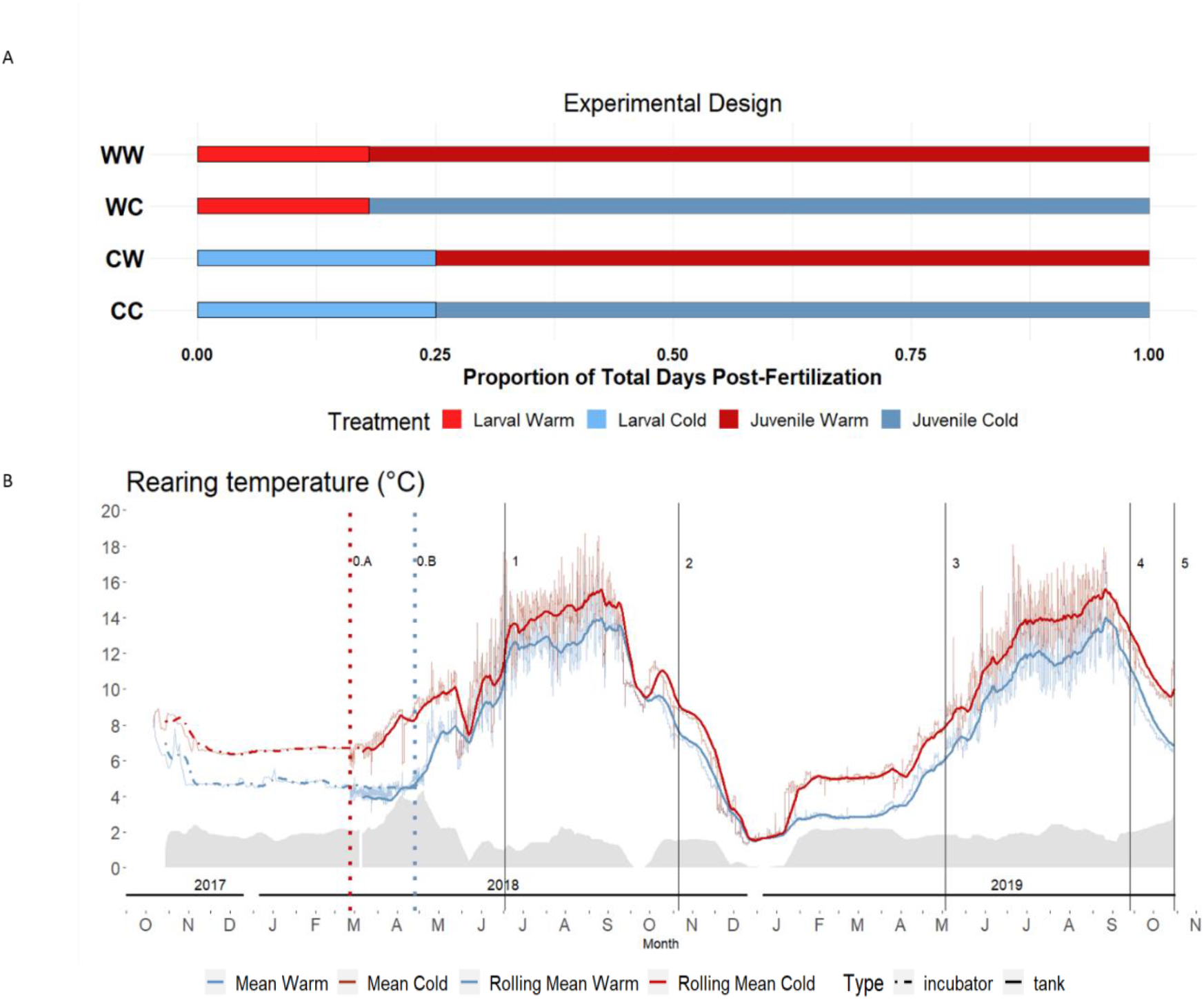
**A)** Experimental design for temperature treatments for larval and juvenile Atlantic salmon (red = warm temperature, blue = cold temperature). Each temperature group of individuals was split into two combinations as outlined above (WW, WC, CW, CC) presented as the proportion of total days post-fertilization across the duration of the experiment. **B)** Temperature curve for the larval and juvenile phases of the experiment with the warm temperature treatment water in red and the cold temperature treatment water in blue. 0.A and 0.B indicate the timing of transport of juveniles to Lammi Biological station for the warm and cold larval treatment individuals, respectively. 1, 2 and 3 indicate the times of routine measurements for length and mass of 464-580 individuals at the Summer0, Autumn0 and Spring1 time points, respectively. 4 and 5 indicate the final two time points with routine measurements of length and mass of 464 and ~1205 individuals, respectively, and maturation status checking in males.

## Statistical Analysis

### Sexual maturation

We fitted a generalized animal model with probit-link function to maturation status at age 2 years (coded as binaries) using Bayesian Markov chain Monte Carlo (MCMC) simulations implemented in MCMCglmm v. 2.32 (Hadfield 2010). We wanted to test whether maturation rates were affected by the larval and juvenile temperature treatments, their interaction, the maturation locus (*vgll3*), and whether the maturation locus (*vgll3*) effects interacted with the larval or juvenile temperature treatment effects or their interaction. We therefore specified the following model to test this and to reflect the mating and experimental designs: *Y = μ + β_1_JuvenileTemperature + β_2_LarvalTemperature + β_3_JuvenileTemperature-By-LarvalTemperature + β_4_Vgll3 + β_5_Vgll3-By-JuvenileTemperature + β_6_Vgll3-By-LarvalTemperature + β_7_Vgll3-By-JuvenileTemperature-By-larvalTemperature + animal-By-JuvenileTemperature + tank-By-JuvenileTemperature + εrror-By-JuvenileTemperature* (1), where *JuvenileTemperature* refers to the juvenile stage rearing temperature, *LarvalTemperature* refers to the larval incubation temperature, and the *β_3_JuvenileTemperature-By-LarvalTemperature* interaction refers to their interaction. The major locus term *Vgll3* refers to a continuous additive effect of *vgll3* genotype (LL = −1, EL or LE = 0, EE= 1, i.e., the additive effect of adding one E allele) and this effect was interacted with all temperature treatment terms. The random terms *animal*, *tank*, and *error*, refer to additive genetic, tank, and residual effects, respectively. The covariance structure for animal was specified as unstructured across the two feeding temperatures and as diagonal for tank and residual effects across the two juvenile rearing temperatures (because their covariance could not be estimated). We fitted variances conditionally on juvenile temperature because we expected larger effects of the juvenile than the larval temperatures. We ran the model with four chains for 1,009,900 iterations each and sampled every 100 iterations. We then ensured that i) sampling convergence was indicated by a scale reduction factor around 1 per chain (Brooks and Gelman 1998), ii) the number of samples to discard (“burn-in“, determined = 100,000) led to consistently reaching a scale reduction factor < 1.1 across chains (Brooks and Gelman 1998), and iii) the thinning per chain resulted in autocorrelations at lag 2 < 0.1 (determined thinning = 500). We also checked for sufficient mixing via MCMC per chain by visually examining the trace plots. These criteria resulted in combined posteriors across chains totaling 7,280 iterations.

### Growth and condition models

We recorded length and mass data at altogether five time points. Because we lethally sampled individuals, we only hold cross-sectional data at the individual level, but obtained longitudinal data (i.e., for several time points) at the biological levels of *vgll3*, sex and family and experimental levels of temperature treatments and tanks. We defined individual body condition as the deviation of the individual mass at the average length as predicted from a regression model of log of mass on log of length. The average length was 10.98 cm so that individual condition is defined as the mass standardized to this length. We fitted general animal models with normally distributed residuals for condition or length records using residual maximum likelihood (REML) as implemented in ASReml-R v. 4.1.0.176 (Butler et al. 2018). We fitted models to the responses of either length or condition that were similar to the model for maturation probability in respect to the two temperature treatments and the major locus, but included additional temporal terms the five time points and SexMat, which characterized a 3-level factor for the combination of sex (female, male) and maturation status (immature, mature, conditional for males because all females were immature).

## Results

### Sexual maturation

No female sampled throughout the study and no male sampled prior to autumn of their second year in fresh water had matured. However, 227 of the 615 males (37%) sampled in autumn of their second year were mature. Results by the generalized mixed model indicated that maturation rates did not differ between the larval temperature treatments reared within each juvenile temperature treatment (Table 1, Figure 2). However, maturation rates did differ between the juvenile temperature treatments with a 2.3 times higher maturation rate in the warm vs. the cold juvenile temperature. Specifically, the back-transformed maturation rate predicted by the generalized mixed model across major locus genotypes and larval rearing temperatures was 0.40 (95% CI: 0.29-0.52) in the warm juvenile rearing treatment and 0.17 (95% CI: 0.10-0.27) in the cold juvenile rearing treatment (warm-cold contrast: 0.23; 0.13-0.32).

**Table 1.**
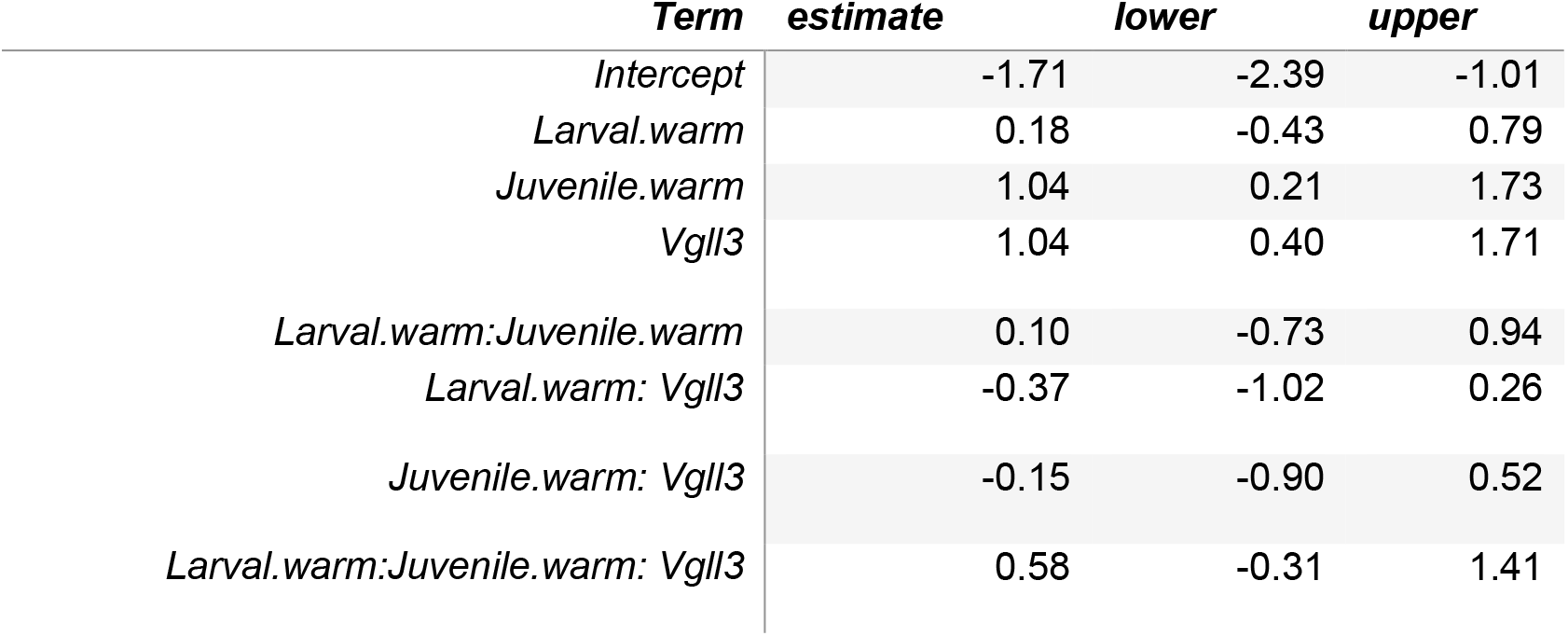
Model coefficient estimates with lower and upper 95% credible intervals for the response of male maturation status (mature, immature). Estimates are on the probit scale.

**Figure 2.**
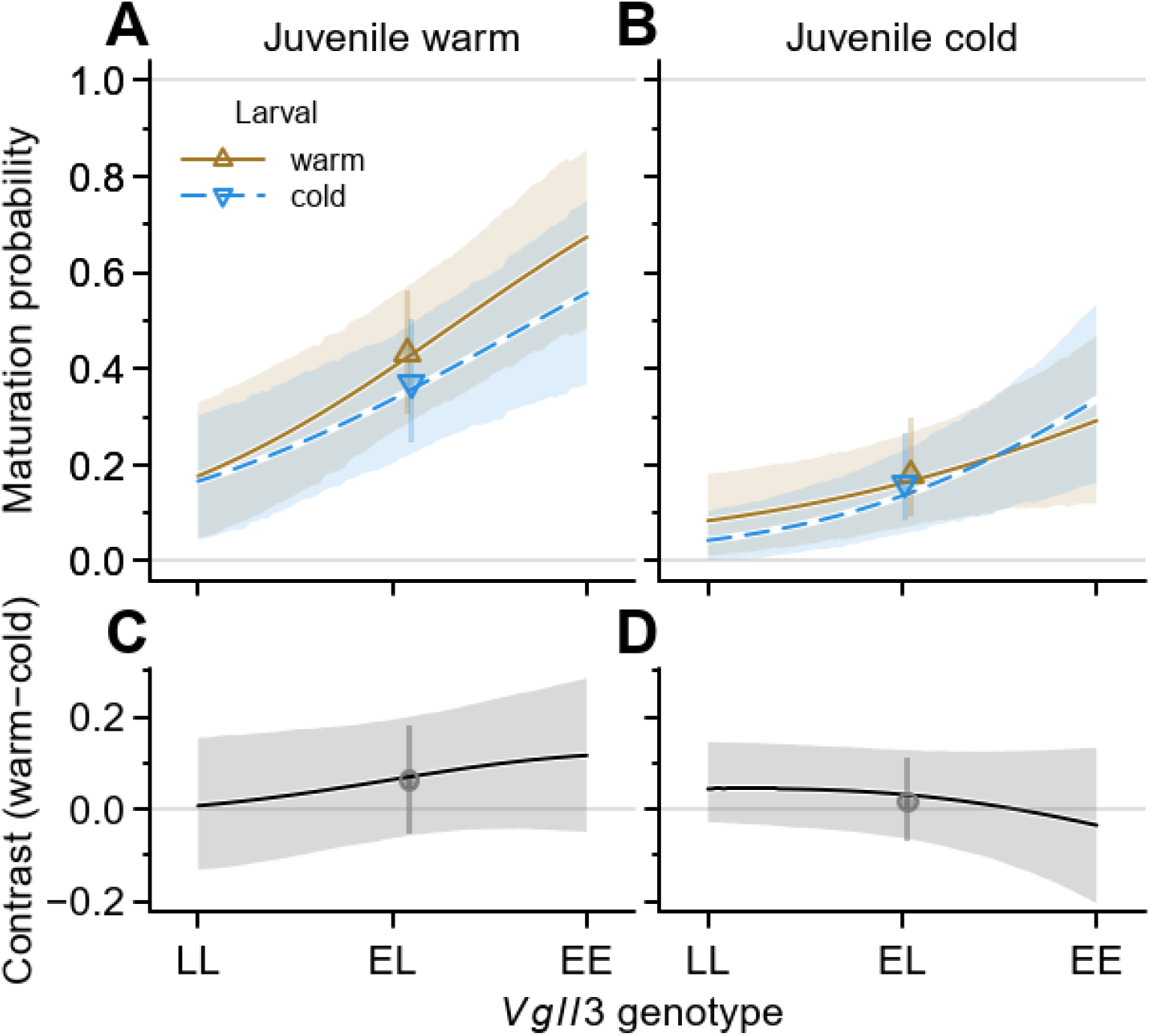
Model predicted, back-transformed male maturation probability for each *vgll3* genotype, and for the overall mean across genotypes, at either a warm (A) or 2°C colder (B) juvenile temperature. The predicted average maturation rates are depicted by larval-temperature-specific symbols with 95% credible intervals and have been plotted at each average *vgll3* allele frequency. The predicted additive major locus (*vgll3*) effects are depicted by larval-temperature-specific lines with 95% credible bands. The corresponding larval-temperature contrasts for both the means and the additive major locus effects are shown in the lower panels (C, D).

The major locus (*vgll3*) affected male maturation rate according to expectations, i.e., adding one or two E alleles dramatically increased the probability to mature from 0.06 to 0.15 and 0.31 in the cold juvenile treatment and from 0.17 to 0.37 and 0.62 in the warm juvenile treatment, respectively (Figure 2, Table 1). In contrast to the average maturation rate, the additive major locus effect did not differ significantly between the larval or juvenile temperatures, or their interaction (Figure 2, Table 1 - model coefficients). In other words, juvenile, but not larval rearing temperature, significantly affected the overall maturation rate and the major locus effect on maturation probability remained consistent regardless of larval temperature rearing treatment.

### *vgll3* genotype effects on condition and length

By predicting results based on a general animal model, it appeared evident that body condition changes between seasons were stronger in *vgll3*\*LL individuals than in *vgll3*\*EE individuals in both sexes: *vgll3*LL* individuals had lower body condition than vgll3*EE individuals in the spring prior to the breeding season, but higher body condition in the autumn, and *vgll3*EL* individuals fell in between (Figure 3 A). The estimates of *vgll3* additive effects for condition and length enabled a formal assessment of these time-specific *vgll3* effects on body condition and the temporal changes of these effects during our experiment. This assessment indicated that body condition is generally affected by *vgll3* in both sexes (Table 2) with the *vgll3* effect during the spring receiving the strongest statistical support, although statistical support was not given once accounted for the false discovery rate (Table 3). However, the change of the *vgll3* effect on condition across time was significant (Table 4). The spring timepoint also had the strongest *vgll3* additive effect contrast between all three other seasons measured in the experiment (Table 4). Thus, even though there was only limited statistical support for time-specific differences in condition among *vgll3* genotypes, the difference in condition change across seasons between *vgll3* genotypes received sufficient statistical support (Table 4).

**Figure 3.**
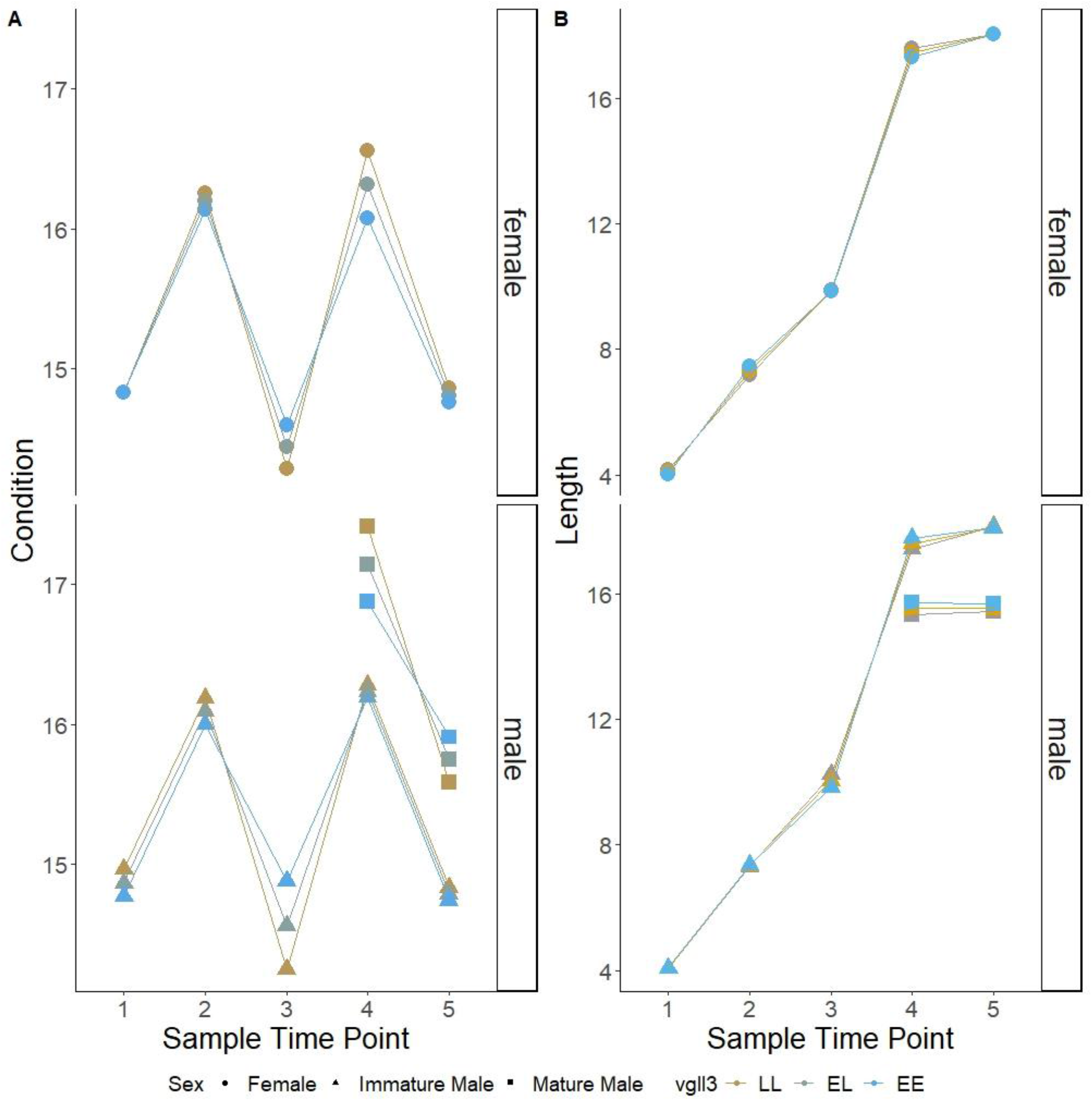
Model-predicted body condition and length values of the large effect maturation locus (*vgll3*) in Atlantic salmon juveniles across 1.5-years (N = 3177). Sample time point numbers 1, 2, 3, 4 and 5 represent 1 – Summer age 0, 2 – Autumn age 0, 3 - Spring age 1, 4 - Autumn age 1 and 5 – Autumn age 1, respectively. The colors represent *vgll3* genotypes (gold = LL, gray = EL, blue = EE) and the shapes represent sex/maturation status (circle = female, triangle = immature male, square = mature male). All females were immature.

**Table 2:** *F*-test results based on the mixed model for body condition.

|  | *DF* | *DDF* | *F* | *P* |
| --- | --- | --- | --- | --- |
| *Intercept* | 1 | 18.3 | 0.01 | 0.9398 |
| *Vgll3* | 1 | 16.1 | 0.22 | 0.6434 |
| *JuvenileTemp* | 1 | 17 | 0.18 | 0.6728 |
| *LarvalTemp* | 1 | 20.9 | 0.72 | 0.4066 |
| *TimePoint* | 4 | 29.5 | 172.3 | <0.001 |
| *Sex.MAT* | 2 | 2015.2 | 150.6 | <0.001 |
| *Vgll3:JuvenileTemp* | 1 | 23.3 | 0.37 | 0.5469 |
| *Vgll3:LarvalTemp* | 1 | 2795.5 | 2.36 | 0.1248 |
| *JuvenileTemp:LarvalTemp* | 1 | 12.4 | 0.00 | 0.9819 |
| *Vgll3:TimePoint* | 4 | 1139.8 | 5.84 | 0.0001 |
| *JuvenileTemp:TimePoint* | 4 | 19.7 | 11.95 | <0.001 |
| *LarvalTemp:TimePoint* | 4 | 29.8 | 0.31 | 0.8703 |
| *Vgll3:Sex.MAT* | 2 | 1960.7 | 2.38 | 0.0929 |
| *JuvenileTemp:Sex.MAT* | 2 | 2023.5 | 0.06 | 0.9372 |
| *LarvalTemp:Sex.MAT* | 2 | 1993.9 | 4.86 | 0.0079 |
| *TimePoint:Sex.MAT* | 5 | 1179.7 | 1.52 | 0.1792 |
| *Vgll3:Juvenile:LarvalTemp* | 1 | 2562.3 | 1.13 | 0.2877 |
| *Vgll3:Juvenile:TimePoint* | 4 | 1123.4 | 0.66 | 0.6176 |
| *Vgll3:LarvalTemp:TimePoint* | 4 | 1133.9 | 1.66 | 0.1559 |
| *Juevenile:LarvalTemp:TimePoint* | 4 | 19.9 | 1.45 | 0.2549 |
| *Vgll3:JuvenileTemp:Sex.MAT* | 2 | 2006.2 | 0.44 | 0.6456 |
| *Vgll3:LarvalTemp:Sex.MAT* | 2 | 1929.9 | 2.89 | 0.0557 |
| *JuvenileTemp:LarvalTemp:Sex.MAT* | 2 | 2015.8 | 6.62 | 0.0014 |
| *Vgll3:TimePoint:Sex.MAT* | 5 | 1157.9 | 0.68 | 0.6395 |
| *JuvenileTemp:TimePoint:Sex.MAT* | 5 | 1157.8 | 0.46 | 0.8063 |
| *LarvalTemp:TimePoint:Sex.MAT* | 5 | 1172.4 | 1.85 | 0.1011 |
| *Vgll3:JuvenileTemp:LarvalTemp:TimePoint* | 4 | 1119.1 | 1.17 | 0.3215 |
| *Vgll3:JuvenileTemp:LarvalTemp:Sex.MAT* | 2 | 1990.8 | 0.98 | 0.3739 |
| *Vgll3:JuvenileTemp:TimePoint:Sex.MAT* | 5 | 1109.1 | 1.39 | 0.2253 |
| *Vgll3:LarvalTemp:TimePoint:Sex.MAT* | 5 | 1150 | 1.23 | 0.2908 |
| *Juvenile:LarvalTemp:TimePoint:Sex.MAT* | 5 | 1139.9 | 1.26 | 0.2804 |
| *Vgll3:JuvenileTemp:LarvalTemp:TimePoint:Sex.MAT* | 5 | 1082.2 | 1.77 | 0.1152 |

**Table 3.** Time-point specific estimates of the *vgll3* additive effect (effect of adding one E allele) on body condition.

| *TimePoint* | *diff* | *sed* | *t* | *p* | *fdr* |
| --- | --- | --- | --- | --- | --- |
| 1Summer0 | 0.00442 | 0.006263 | 0.72 | 0.490474 | 0.613 |
| 2Autumn0 | 0.006389 | 0.006009 | 1.06 | 0.303397 | 0.506 |
| 3Spring1 | -0.01447 | 0.006415 | -2.26 | **0.038369** | 0.192 |
| 4Autumn1 | 0.010562 | 0.006085 | 1.74 | 0.101708 | 0.254 |
| 5Autumn1 | 0.002602 | 0.005175 | 0.50 | 0.621928 | 0.622 |

**Table 4.**
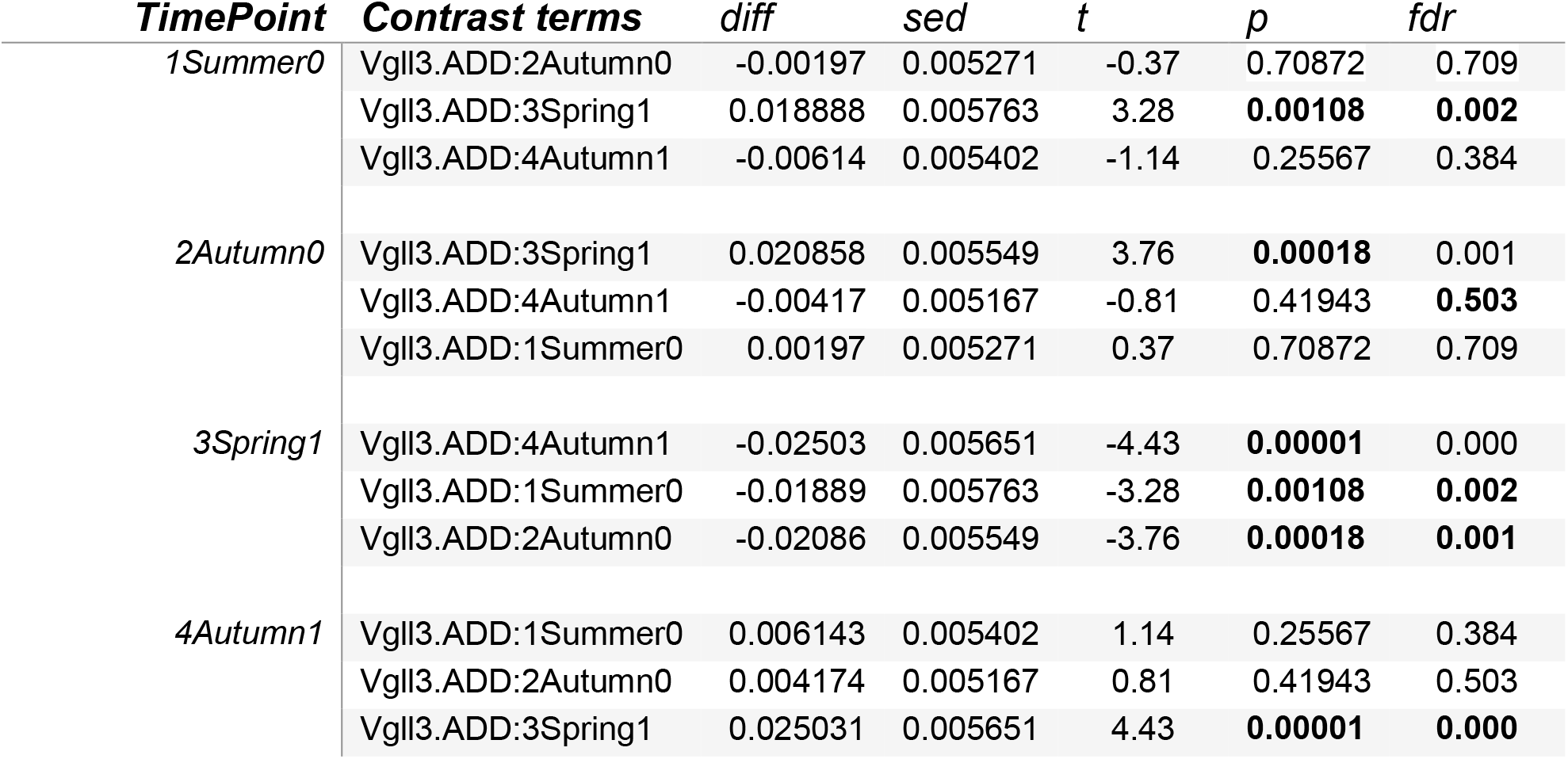
Between-time point contrasts of the *vgll3* additive effect (effect of adding one E allele) on body condition.

In contrast to condition, length was not affected by *vgll3* (Table 5). However, mature *vgll3*\*LL males in CC were fatter but shorter than *vgll3*\*EE males, whereas mature *vgll3*\*LL males in WW were thinner but longer than *vgll3*\*EE males, but only at the first time point in autumn as shown above in Figure 3.

**Table 5:** *F*-test results based on the mixed model for length.

|  | *DF* | *DDF* | *F* | *P* |
| --- | --- | --- | --- | --- |
| (Intercept) | 1 | 16.7 | 13190 | <0.001 |
| Vgll3 | 1 | 16.8 | 0.8 | 0.380 |
| JuvenileTemp | 1 | 23.5 | 224.3 | <0.001 |
| LarvalTemp | 1 | 22 | 42 | <0.001 |
| TimePoint | 4 | 28.1 | 12230 | <0.001 |
| Sex.MAT | 2 | 2134.5 | 108.6 | <0.001 |
| Vgll3: Juvenile Temp | 1 | 19.9 | 2.4 | 0.1396 |
| Vgll3:LarvalTemp | 1 | 2329.6 | 2.5 | 0.1113 |
| *JuvenileTemp:LarvalTemp* | 1 | 31.3 | 0.7 | 0.3954 |
| *Vgll3:TimePoint* | 4 | 1140.1 | 1.8 | 0.119 |
| *Juvenile Temp:TimePoint* | 4 | 22.6 | 14.7 | <0.001 |
| *Larval Temp:TimePoint* | 4 | 28.4 | 7.9 | 0.0002 |
| *Vgll3:Sex.MAT* | 2 | 2103.9 | 0.2 | 0.8001 |
| *JuvenileTemp:Sex.MAT* | 2 | 1942.4 | 3.2 | 0.0409 |
| *LarvalTemp:Sex.MAT* | 2 | 2118.8 | 0.2 | 0.8277 |
| *TimePoint:Sex.MAT* | 5 | 1238.1 | 1.1 | 0.3331 |
| *Vgll3: JuvenileTemp:LarvalTemp* | 1 | 2099.5 | 0.2 | 0.6826 |
| *Vgll3: JuvenileTemp:TimePoint* | 4 | 1108.7 | 0.7 | 0.5683 |
| *Vgll3: LarvalTemp:TimePoint* | 4 | 1136.5 | 2.1 | 0.0759 |
| *JuvenileTemp: Larval Temp:TimePoint* | 4 | 22.9 | 0.4 | 0.7956 |
| *Vgll3: Juvenile Temp:Sex.MAT* | 2 | 1968.1 | 1.3 | 0.275 |
| *Vgll3: Larval Temp:Sex.MAT* | 2 | 2083.3 | 0.7 | 0.4977 |
| *Juvenile Temp: Larval Temp:Sex.MAT* | 2 | 1935.4 | 4 | 0.0179 |
| *Vgll3:TimePoint:Sex.MAT* | 5 | 1224.9 | 2 | 0.0812 |
| *Juvenile Temp:TimePoint:Sex.MAT* | 5 | 1202.9 | 1.9 | 0.0848 |
| *Larval Temp:TimePoint:Sex.MAT* | 5 | 1231.9 | 1.4 | 0.2134 |
| *Vgll3: Juvenile Temp:LarvalTemp:TimePoint* | 4 | 1104 | 0.8 | 0.5358 |
| *Vgll3: JuvenileTemp:Larval Temp:Sex.MAT* | 2 | 1943.6 | 1.3 | 0.2761 |
| *Vgll3: JuvenileTemp:TimePoint:Sex.MAT* | 5 | 1163.1 | 2 | 0.0782 |
| *Vgll3: LarvalTemp:TimePoint:Sex.MAT* | 5 | 1214.9 | 1.7 | 0.1219 |
| *JuvenileTemp: Larval Temp:TimePoint:Sex.MAT* | 5 | 1194.1 | 1.5 | 0.1883 |
| *Vgll3: JuvenileTemp:LarvalTemp:TimePoint:Sex.MAT* | 5 | 1143.9 | 0.9 | 0.4868 |

### Larval temperature effects on body condition, length, and interaction with *vgll3* genotypes

Body condition showed only one significant (FDR < 0.05) effect on means between larval temperatures (ignoring *vgll3*): in mature males at the last time point and at the warm feeding temperature with WW males being 0.6 g (0.3-0.9 g) smaller for the same length as CW males. With length, there were several significant (FDR < 0.05) effects on means between larval temperatures. However, no differences for any *vgll3* effects with larval temperatures were detected for either condition or length, only the abovementioned *vgll3* effects on condition change.

## Discussion

It has previously been shown that temperature experienced at an early point in life can have lasting effects in later life stages (Macqueen et al. 2008; Nord and Nilsson 2016; While et al. 2018; reviewed in Jonsson and Jonsson 2019) We aimed to experimentally test for the occurrence of such effects during the freshwater phase in Atlantic salmon as water temperature has been earlier shown known to influence behavior, food availability and metabolic rate (Morash et al. 2021; Jutfelt et al. 2021). We present two key findings that contribute to our understanding how temperature differences experienced during early life affect (life-history) traits later in life. The first finding that a 2⁰C difference in rearing temperature during the larval phase, the 4.5- to 6-month period from fertilization to first exogenous feeding, did not significantly affect maturation rate, is of relevance for considering the fate of wild populations experiencing environmental change, but also to aquaculture production where early sexual maturation is an undesired event. Juveniles of most wild Atlantic salmon populations spend at least two, and sometimes more, years in riverine environments where water temperatures can vary considerably between larval and juvenile phases. Our results suggest that populations may be resilient to temperature differences of up to 2⁰C during the larval phase when considering future effects on maturation age, at least at the age of two years as mature parr as studied here. Our second finding of relevance is that differing larval or juvenile temperature did not appear to alter the *vgll3* locus effects on maturation, nor on growth, length, or body condition. Interestingly, no differences for any sex in any feeding temperature exist thereafter which meant the individuals in the colder temperature during larval phase had full growth compensation during the length of the experiment.

This second finding of no larval or juvenile temperature effect with *vgll3* leads us to expect a temperature consistency in *vgll3* effects on maturation and other affected traits, and thus also to a *vgll3*-effect consistency in response to either natural or artificial selection. This is an important finding with respect to modelling the effects of temperature change across thermal environments as the predicted effects of *vgll3* on maturation probability can be assumed stable across temperatures. That said, it should be noted that a relatively narrow range of temperatures were explored here, so future research across a broader temperature range may be useful. Lastly, the overall higher maturation probability in the warmer juvenile treatment observed in controlled conditions in our study is consistent with previous reports in wild populations (Martinez et al. 2000) and controlled conditions (Rowe and Thorpe 1990a; Åsheim et al. 2022) showing higher rates of early maturation in certain populations with warmer temperatures (Jonsson and Jonsson 2013). We found *vgll3* genotype was correlated with higher maturation probability with each addition of a *vgll3*\*E allele increasing the observed maturation probability. This was also found in a recent study by Debes et al. (2021) which found the additive effect of one *vgll3*\*E allele probability estimate to be 0.94 compared to 1.04 in our study. In addition, this is the first time to our knowledge of *vgll3* interactions with different larval rearing temperatures being reported and with *vgll3* showing no effect with the warm and cold larval temperature treatments. The observed seasonal changes in body condition were as expected, with body condition being highest in the autumn, following the period of highest food consumption rate in the summer, and lowest in the spring following the over-wintering period at cold temperatures (Figure 3A). However, an unexpected, but nevertheless noteworthy, finding of this study was that *vgll3* genotype affected the level of seasonal change in body condition in both sexes whereby *vgll3*LL* individuals had lower body condition than *vgll3*EE* individuals in the spring prior to the breeding season, but then higher body condition in the autumn. These *vgll3* genotype specific seasonal changes in body condition over the 1.5-year study period add further nuance to previous studies that suggested a role of *vgll3* in the control of resource allocation (Debes et al. 2021; Halperin et al. 2013). *Vgll3* effects on body condition may express as effects on condition change and, and thus may or may not express as an effect on average condition at a given time point. Our findings suggest that the general assumption that individuals with higher body condition are more likely to mature earlier due to having higher lipid reserves (Andersson et al., 2018; Good & Davidson, 2016; Roff, 2002; Rowe et al., 1991; Stearns, 1992; Taranger et al., 2010; Wells et al., 2017) may be too simplistic. Rather, backing up previous statements that it may be that having an adequate storage of energy at critical life history timepoints, which for salmon is thought to be in the spring prior to maturation, is key (Rowe et al, 1991). This is indeed the timepoint at which juveniles carrying the *vgll3*EE* genotype exhibited higher body condition than individuals carrying other *vgll3* genotypes (Figure 3a), even though body condition was recorded at its lowest point for all genotypes of the five timepoints measured.

Our observation that the body condition of *vgll3*EE* individuals was more stable across seasons than that of *vgll3*LL* individuals is in line with recent studies investigating links between *vgll3* genotypes and other juvenile phenotypes including aggressive behavior and aerobic scope, both of which could perceivably have an effect on condition factor in Atlantic salmon. It was earlier found that *vgll3*LL* juveniles were more aggressive compared to *vgll3*EE* individuals (Bangura et al. 2022). Such a behavioral difference could result in *vgll3*LL* individuals allocating energy for aggressive behavior, which otherwise could have been allocated to lipid storage. Considering aerobic scope, it was found that *vgll3*EE* individuals had higher aerobic scope than *vgll3*LL* individuals. Thus, superior resource acquisition or assimilation via higher aerobic scope was suggested as a potential mechanism by which an increased condition factor in *vgll3*EE* individuals could be achieved (Prokkola et al. 2022). Our finding here suggest that these qualities may be particularly important during the winter months, when *vgll3*LL* individuals lost body condition much faster than *vgll3*EE* individuals, the result being that *vgll3*EE* individuals had higher condition factor at the critical point in the spring when physiological processes related to maturation are being determined.

One potential caveat for interpreting these results is the strong decline in body condition observed between the last two measuring time points just several weeks apart with mature individuals included in this calculation. One potential explanation for this is the prolonged sampling from time point 4 to 5, resulting from the large number of individuals being measured, may have resulted in the longer period of fasting resulting in lower condition in individual sampled during the fifth time point. This pattern could also simply be due to routine sampling involving dissection and growth measurements over a prolonged period of time e.g. one month, which is indicated by a decreasing *vgll3*\*LL mature male length with time (Figure 3B, male panel), which is unexpected over such as short time period, but could be explained by random sampling from tanks. Alternatively, males may need different cues (high condition vs. high length) to become mature in the different temperature environments of WW vs. CC as we see the average length to be the same at the first time point for both sexes and in both feeding temperatures. These results may simply reflect the later initiated feeding of the cold incubated fish.

To conclude, our study provides details of how genetic (the *vgll3* locus) and environmental (seasonal temperature) effects contribute to maturation probability, with seasonal body condition being a central phenotype. Importantly, the seasonal context in which condition factor is measured needs to be considered when interpreting results, as the relative condition factors of individuals with differing *vgll3* genotypes was completely reversed in autumn vs. spring. Future work to better understand energy allocation processes e.g., via lipidomics or functional genomics could help to shed more light on the mechanisms by which the large-effect *vgll3* locus influences maturation and exploring a broader range of temperature differences could aide understanding the absence of an effect of larval temperature on maturation.

## Acknowledgements

We acknowledge Ksenia Zueva, Marion Sinclair-Waters, Nico Lorenzen, Spiros Papakostas, and Victoria Pritchard for their assistance in getting fish gametes in Laukaa hatchery; Suvi Ikonen, Anna Toikkanen, Shadi Jansouz, Dorian Jagusch, Andres Salgado, Fin Morrison, Ike Van Gestel, Paul Bangura, and Petra Lijeström for their help in Lammi with husbandry and fish sampling and Outi Ovaskainen; Miakel Kyriacou, Oona Mehtälä, Oliver Andersson, Pirta Palola, Valeria Valanne, and Jacqueline Moustakas-Verho for their help during sampling events; Annukka Ruokolainen, Noora Parre, Shadi Jansouz and Seija Tillanen for their help in the genetics lab and Eirik Åsheim for his help with manuscript discussions and data wrangling.

## Notes

**Funding** This work was funded by the University of Helsinki LUOVA Doctoral School, Fulbright Finland, Academy of Finland [grant numbers 314254, 314255, 327255] and the European Research Council under the European Union’s Horizon 2020 research and innovation program [grant agreement number 742312].

**Conflict of Interest Statement** The authors declare no conflicts of interest.

### Competing Interest Statement

The authors have declared no competing interest.

